# IL4I1 binds to TMPRSS13 and competes with SARS-Cov2 Spike

**DOI:** 10.1101/2021.12.23.473686

**Authors:** Jérôme Gatineau, Charlotte Nidercorne, Aurélie Dupont, Marie-Line Puiffe, José L Cohen, Valérie Molinier-Frenkel, Florence Niedergang, Flavia Castellano

**Author notes:** Univ Paris Est Creteil, INSERM, IMRB, F-94010 Creteil, France. Corresponding authors: Flavia Castellano*, INSERM U955, IMRB eq J-Biot, Hôpital Henri Mondor, 1 rue Eiffel, F-94010 Créteil cedex, France; *, Florence Niedergang* Institut Cochin, 22 Rue Méchain, 75014 Paris, Valérie Molinier-Frenkel*, INSERM U955, IMRB eq I-Biot, Hôpital Henri Mondor, 1 rue Eiffel, F-94010 Créteil cedex, France.

## Abstract

The secreted enzyme interleukin four-induced gene 1 (IL4I1) is involved in the negative control of the adaptive immune response. IL4I1 expression in human cancer is frequent and correlates with poor survival and resistance to immunotherapy. Nevertheless, its mechanism of action remains partially unknown. Here, we identified transmembrane serine protease 13 (TMPRSS13) as an immune cell-expressed surface protein that binds IL4I1. TMPRSS13 is a paralog of TMPRSS2, whose protease activity participates in the cleavage of SARS-Cov2 Spike protein and facilitates virus induced-membrane fusion. We show that TMPRSS13 is expressed by human lymphocytes, monocytes and monocyte-derived macrophages, can cleave the Spike protein and allow Sars-Cov2 Spike pseudotyped virus entry into cells. We identify regions of homology between IL4I1 and Spike and demonstrate competition between the two proteins for TMPRSS13 binding. These findings may be relevant for both interfering with SARS-Cov2 infection and limiting IL4I1-dependent immunosuppressive activity in cancer.

**One-Sentence Summary:** Through binding to its newly identified receptor TMPRSS13, the enzyme IL4I1 interferes with SARS-Cov2 Spike cleavage thereby blocking viral entry into host cells.

## Introduction

Negative control of the immune response is achieved through multiple mechanisms, amongst which expression of the enzyme interleukin four-induced gene 1 (IL4I1). IL4I1 is a secreted L-amino acid oxidase that catabolizes the essential amino acid phenylalanine and, to a lesser extent tryptophan and arginine, into the corresponding α-keto acids, H2O2, and NH3 (*1*). Antigen-presenting cells are the main producers of IL4I1 under physiological conditions (*2, 3*). IL4I1 regulates B and T-cell activation and proliferation (*3*–*5*) and facilitates the differentiation of FoxP3^+^ regulatory T cells from naïve CD4^+^ T cells (*6*). In the context of cancer, IL4I1 is expressed by infiltrating macrophages and occasionally by the tumor cells (*7*). In mouse tumor models, IL4I1 facilitates escape from the immune response (*8*) and in humans, its local expression correlates with decreased survival and a pejorative outcome (*9–11*).

The mechanisms of action of IL4I1 are only partially known. The rapid IL4I1-mediated inhibitory effect on TCR signaling, within minutes from T-cell activation, its focal secretion into the synaptic cleft, and its detection on the T-cell membrane (*5*), suggest the existence of an IL4I1 receptor.

Patients with severe Coronavirus disease 19 (COVID-19), caused by severe acute respiratory syndrome Coronavirus 2 (SARS-Cov2), present profound dysregulation of the immune response, characterized by hyper-inflammation (*12*) and lymphopenia (*13*). Virus entry into cells depends on binding of the spike protein to cellular receptor angiotensin converting enzyme 2 (ACE2) and on its priming by host cell transmembrane serine protease (TMPRSS) 2 (*14, 15*). Spike cleavage leads to the liberation of an S1 subunit that contains the receptor-binding domain, while exposing the S2 fragment, which allows fusion of the viral envelop with the cell membrane. The TMPRSS proteins are type II transmembrane serine proteases. Activation of the zymogen requires the cleavage of the C-terminal extracellular part of the protein, which remains attached to the cell surface through intramolecular disulfide bonds (*16*). Other proteases of this family have been shown to activate spike-pseudotyped virus (*17*).

Here, we searched for an IL4I1 receptor at the surface of immune cells and identified TMPRSS13, a poorly known paralogue of TMPRSS2. This protease can substitute for TMPRSS2, allowing infection of human monocytes-derived macrophages (hMDM) by Spike-pseudotyped virus (LVS). Most interestingly, IL4I1 and SARS-Cov2 spike share regions of homology and can compete for TMPRSS13 binding.

## Results

### Identification of TMPRSS13 as an IL4I1 receptor

We used the ligand receptor capture (LRC)-TriCEPS® technique (*18*) to identify the protein(s) responsible for IL4I1 binding to the surface of T lymphocytes. We conjugated the TriCEPS reagent with a positive control (transferrin for Jurkat T cells or anti-CD28 antibodies for primary T lymphocytes), a negative control (glycine), or recombinant human IL4I1. We next incubated these conjugates at 4°C with the Jurkat cells (**fig. 1A)** or sorted primary CD3^+^ lymphocytes (**fig. 1B)** and revealed the cell-surface-bound conjugates by flow cytometry (FCM) after TriCEPS staining (gating strategy **fig. S1**). No TriCEPS-glycine bound to Jurkat cells, whereas almost all the cells were labeled with TriCEPS-transferrin. In accordance with our previous observations (*5*), TriCEPS-IL4I1 bound to Jurkat cells. The MFI of TriCEPS-IL4I1 was lower than that of TriCEPS-transferrin (432 ± 110.00 and 3888.3 ± 1439.2 respectively), indicating a lower level of expression of the IL4I1 receptor than that of the transferrin receptor. When we tested primary CD3^+^ T lymphocytes, ~90% percent of the CD3^+^ population bound the TriCEPS-CD28 conjugate (**fig. 1B-F**). A fraction of both CD4^+^ and CD4^-^ CD3^+^ cells (9.66% ± 0.08 and 4.22% ± 0.29 respectively) bound TriCEPS-IL4I1, confirming a specific surface interaction of IL4I1 with T lymphocytes. We further confirmed IL4I1 binding using TAMRA-labeled TriCEPS and immunofluorescence microscopy (IF) (**fig. 1G)**. Conjugates were visible at the cell surface for both transferrin and IL4I1, whereas we detected no signal with glycine. In accordance with the FCM results, the labeling was more intense for transferrin than for IL4I1. Thus, IL4I1 binds specifically to T lymphocytes.

**Figure 1.**
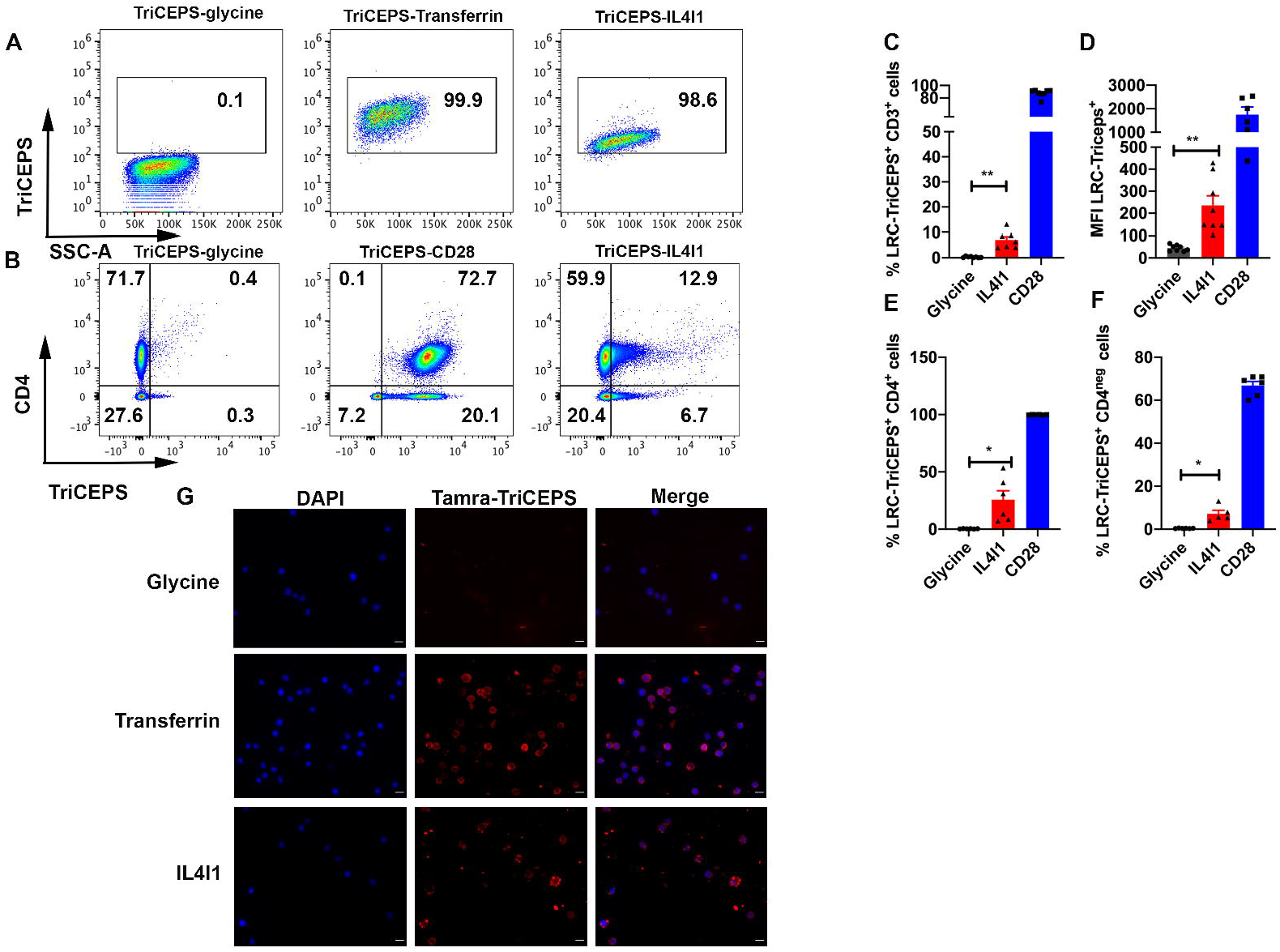
IL4I1 binds to Jurkat cells and human primary T lymphocytes. (**A**) Jurkat cells were incubated with TriCEPS-glycine, TriCEPS-transferrin, or TriCEPS-IL4I1 at 4°C for 1 h. (**B**) Purified CD3^+^ primary T cells were similarly treated (TriCEPS-anti-CD28 antibodies were used instead of TriCEPS-transferrin). After binding, cells were labeled with a viability dye and streptavidin to reveal bound TriCEPS-conjugates. For T lymphocytes, cells were also labeled with an anti-CD4 antibody before analysis by FCM. Percentage of TriCEPS-positive CD3 lymphocytes (**C**), mean intensity of fluorescence (MFI) of TriCEPS on CD3^+^ lymphocytes (**D**), percentage of TriCEPS-positive CD4^+^ T cells (**E**), and TriCEPS-positive CD4^-^ T cells (**F**). N= 4 for CD3^+^ cells and N = 3 for CD4^+^. **(G)** Visualization by IF of IL4I1 binding using TAMRA-LCR-TriCEPS. Jurkat cells were incubated with TAMRA-TriCEPS bound to glycine (top), transferrin (middle), or IL4I1 (bottom). Nuclei were labeled with DAPI. N =3. Bar = 10μm.

We next sought to identify the protein(s) implicated in IL4I1 binding. We cross-linked the TriCEPS-IL4I1 conjugate to the Jurkat cell surface by exposing the cells to gentle oxidizing conditions with sodium-metaperiodate. TriCEPS-transferrin was used as a positive control. Cells were then lysed and analyzed by liquid chromatography/mass spectrometry (LC-MS) (**fig. 2 A&B**). Mass spectrometry analysis led to the identification of 163 differentially present proteins. Twenty-one proteins in the samples incubated with TriCEPS-IL4I1, showed a Log2 fold change (FC) of 2 or more, including IL4I1 itself. However, aside from IL4I1, only nine were significantly enriched. Amongst them, the serine transmembrane protease TMPRSS13 showed a FC of 7.2, similar to the enrichment of the IL4I1 bait (Log2 FC of 6.7). We thus focused on this protein for subsequent analysis.

**Figure 2.**
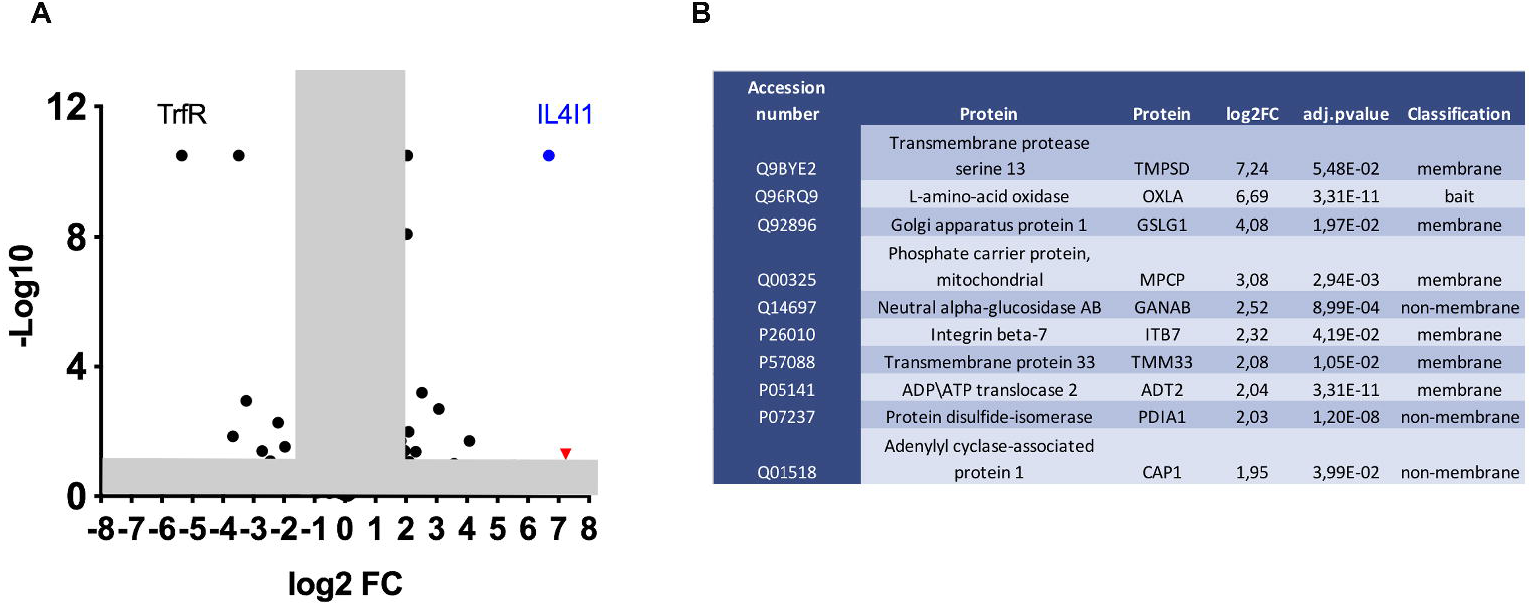
Identification of potential IL4I1 receptor candidates by TriCEPS/LC-MS. Jurkat cells were incubated with TriCEPS-transferrin or TriCEPS-IL4I1 followed by LC/MS. (**A**) Comparative data are shown as a volcano plot, with p < 0.05 and the fold change (log2 FC) > 2 outside the shaded box. Data were derived from three independent technical replicates. TMPRSS13 red dot, IL4I1 blue dot. (**B**) List of the identified proteins by TriCEPS-IL4I1 captures.

We cloned the TMPRSS13 cDNA and stably expressed a DYK-tagged protein in HEK cells (**fig S2**) and chose a strong expressing clone (HEK-T) verified by western blotting (WB) and IF using an anti-tag antibody. The recombinant protein showed an apparent molecular weight of approximately 70 kDa (**fig. S2 A&B**), as already described (*19*). We detected other bands using an anti-TMPRSS13 antibody that recognizes the C-terminal part of the protein on transiently transfected cells (**fig. S2 C**). These bands likely correspond to intermediate products of glycosylation or cleavage of the extracellular C-terminus of the protein, as described by Murray et al. (*19*). The IF labeling of the HEK-T cells with anti-tag (DYK) and with the commercially available anti-TMPRSS13 antibody directed towards the C-terminus of TMPRSS13 were comparable (**fig. S2 E-H**), allowing us to validate this antibody for the study of cells naturally expressing TMPRSS13.

To confirm the interaction of IL4I1 with TMPRSS13 on HEK-T and Jurkat cells, we first performed the proximity ligation assay (PLA). HEK and HEK-T cells were incubated with recombinant IL4I1 and processed for PLA using a rabbit primary antibody against IL4I1 and a mouse antibody directed against the DYK tag of recombinant TMPRSS13. No positive signal was detectable in the experimental controls nor in HEK cells incubated with IL4I1, whereas a strong PLA-positive signal was visible on the HEK-T cells incubated with the protein, indicating proximity between IL4I1 and TMPRSS13 **(fig. 3**). On average, the cells showed 4.1 ± 2.4 dots per cell. A similar signal was detected on Jurkat cells but with fewer spots per cell (1.3 ± 0.4) (**fig. S3**).

**Figure 3.**
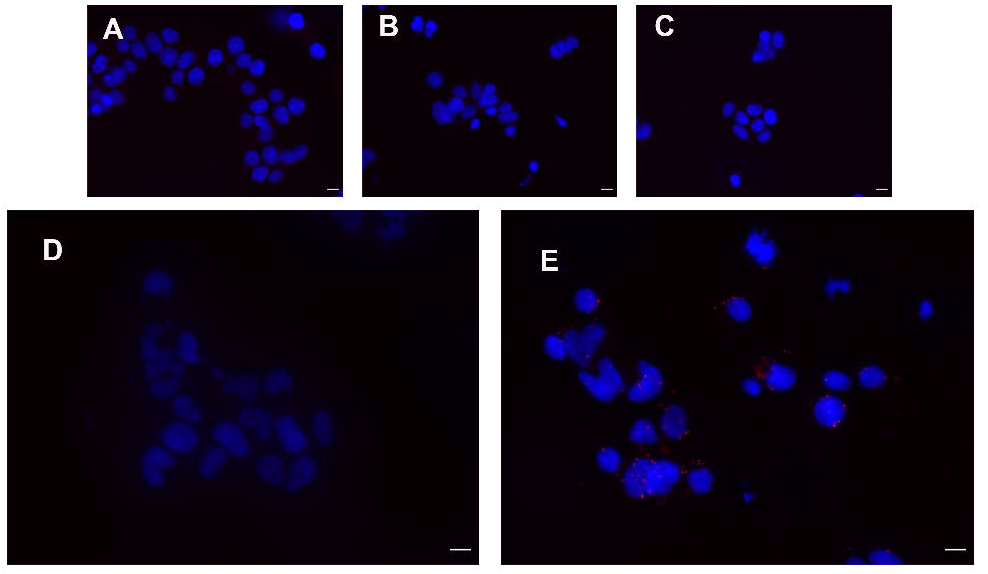
Proximity ligation assay reveals spatial proximity of IL4I1 and TMPRSS13. HEK or HEK-T cells were incubated with recombinant hIL4I1 at 4°C for 1 h and tested by the PLA. Nuclei were stained with DAPI. (**A**) No primary antibodies, (**B**) primary anti-DYK antibody only, (**C**) primary anti-IL4I1 antibody only, (**D**) HEK cells incubated with IL4I1 and primary and secondary antibodies, (**E**) HEK-T cells incubated with IL4I1 and primary and secondary antibodies. N =3. Bar = 10 μM.

As PLA indicates only the proximity of IL4I1 and TMPRSS13, we further validated the interaction by performing co-immunoprecipitation (Co-IP) experiments (**fig. 4**). After incubation with IL4I1, HEK control cells and HEK-T cells were lysed and TMPRSS13 (**fig. 4A**) or IL4I1 **(fig. 4B**) precipitated, respectively. IL4I1 was co-immunoprecipitated with TMPRSS13 (**fig. 4A),** as revealed using either an anti-tag (Myc) or anti-IL4I1 antibody. We observed no precipitation when TMPRSS13 was not expressed. In the reverse Co-IP, we specifically detected TMPRSS13 after precipitation of IL4I1 in HEK-T cells only **(fig. 4B**). Thus, IL4I1 and TMPRSS13 directly interact with each other and TMPRSS13 represents an IL4I1-receptor at the surface of T cells.

**Figure 4.**
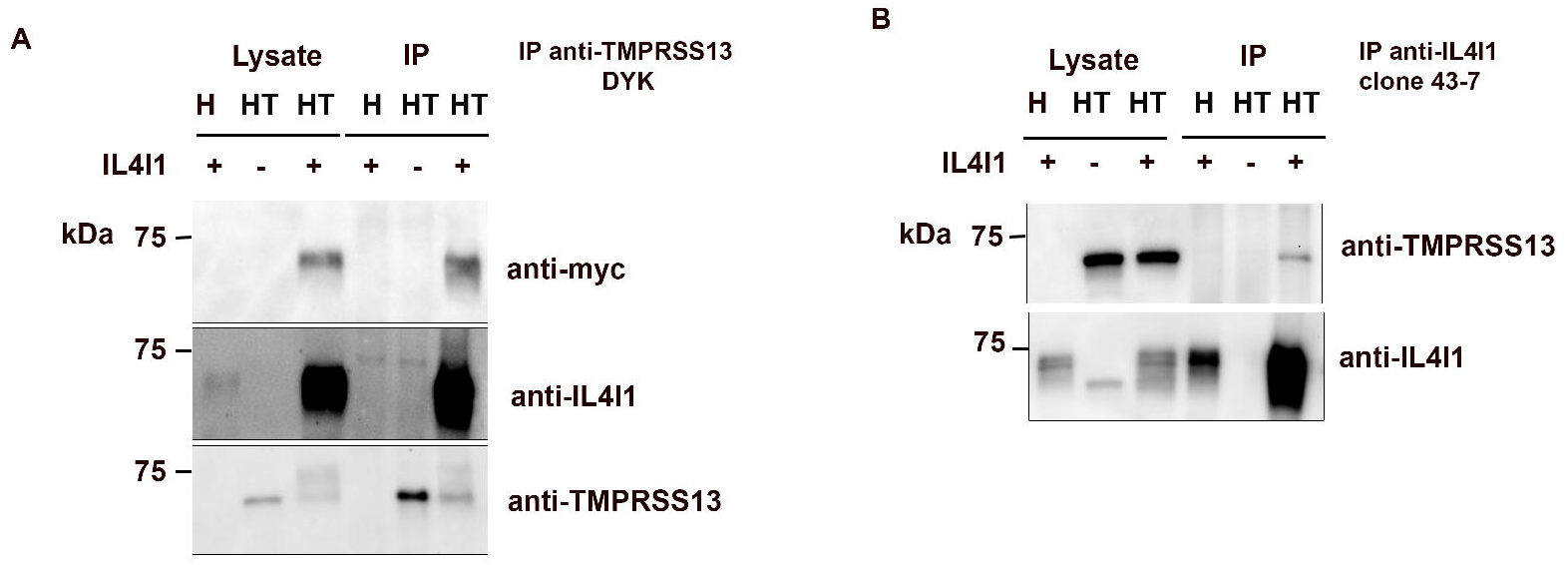
IL4I1 and TMPRSS13 co-immunoprecipitate. Concentrated Myc-tagged IL4I1 was added to wells containing HEK or HEK-T cells and incubated for 1 □h at 4□°C. Cells were lysed and (**A**) immunoprecipitated with anti-DYK antibody (for TMPRSS13) and the precipitates analyzed by WB using the indicated antibodies. (**B**) Reverse co-IP of (A) performed with anti-IL4I1 antibodies. N>3.

### Expression of TMPRSS13 by immune cells

Little information is available concerning physiological TMPRSS13 protein expression, but RT-PCR data indicate that it may be produced by immune cells (*20*). Thus, we verified TMPRSS13 expression by labeling Jurkat cells with the validated anti-TMPRSS13 antibody (**fig. 5**). The Jurkat cells expressed TMPRSS13 at the cell surface, as visualized by IF (**fig. 5A**) and FCM (**fig. 5B**). We next analyzed TMPRSS13 expression by WB on several T-cell lines, including Jurkat cells, and on sorted primary human CD3^+^ and CD20^+^ lymphocytes using the validated antibody, which is directed towards the C-terminus, and two different commercial antibodies targeting N-terminal epitopes (**fig. 5C**). Both antibodies targeting the N-terminal epitopes revealed a protein of approximately 70 kDa in all lysates, corresponding to the molecular weight of the full-length TMPRSS13. However, when the C-terminus-directed antibody was used, the full-length protein was barely visible. By contrast, a strong signal at approximately 30 kDa was revealed in all T-cell lines and primary T cells but was absent from B cells. The molecular weight of this fragment corresponds to that of the C-terminal catalytic domain (*19*). We next analyzed TMPRSS13 expression on human peripheral blood mononuclear cells (PBMCs) by IF (**fig. 5D**). Some T cells were TMPRSS13^+^ and showed fainter signals then larger CD3^-^ cells, possibly monocytes. We refined our analysis by FCM (**fig. 5E&F**) by measuring TMPRSS13 expression in CD4^+^ and CD8^+^ T cell subsets, B cells, monocytes, and NK cells. All showed expression of TMPRSS13, with large differences in the percentage of positive cells between PBMCs from different donors, but with a relatively similar MFI of the corresponding TMPRSS13^+^ population. Based on the MFI, monocytes, B lymphocytes, and NK cells showed the highest level of TMPRSS13 expression. We differentiated macrophages from circulating monocytes and tested the expression of TMPRSS13 by surface IF. The level of expression was variable in these cells showing fine punctate staining at the cell surface and was confirmed by FCM (**fig. 5G)**.

**Figure 5.**
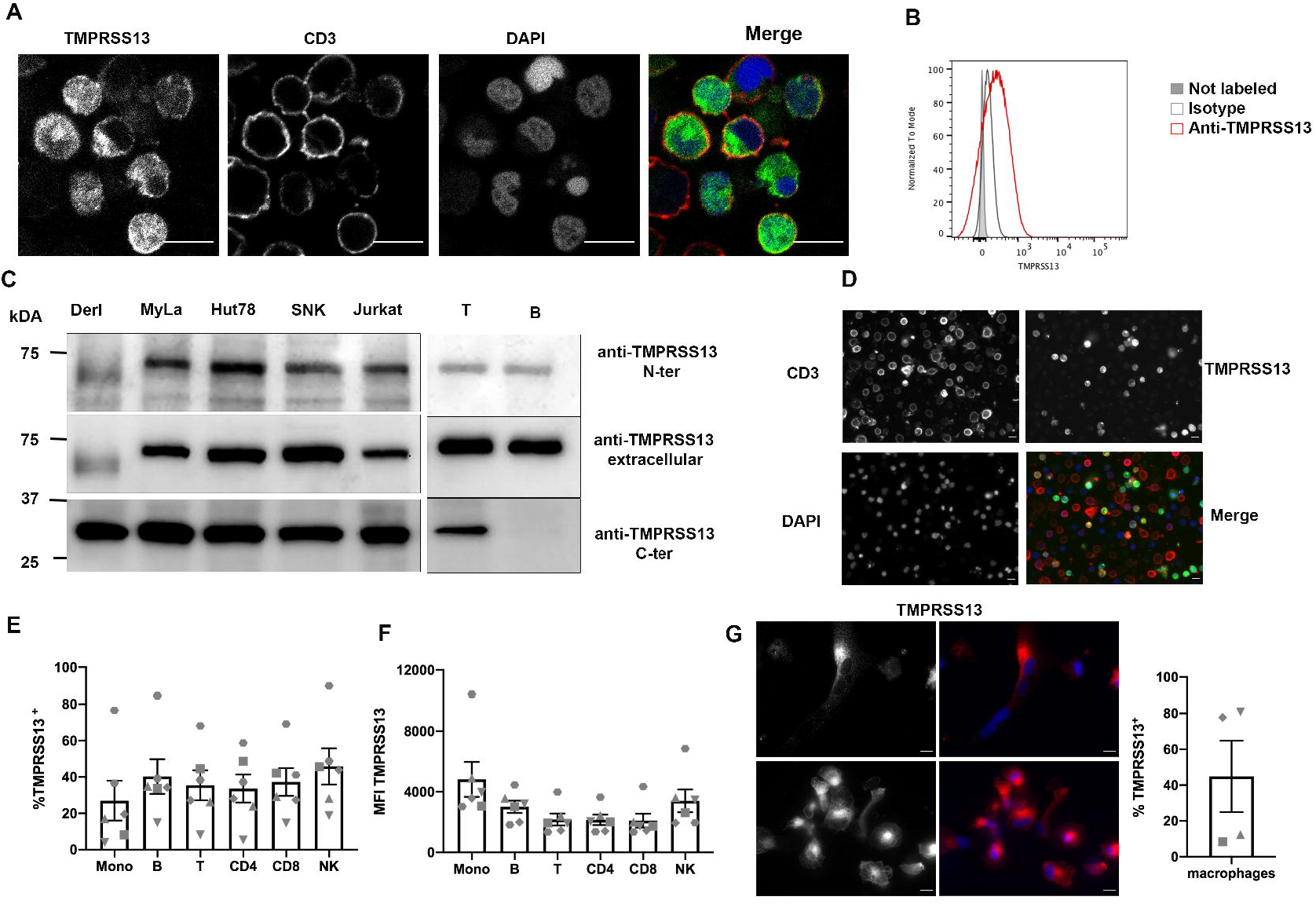
Expression of TMPRSS13 by Jurkat cells and human circulating immune cells. **(A)** Immunofluorescence surface labeling of Jurkat cells with anti-TMPRSS13 (C-ter) (green) and anti-CD3 (red) antibodies. Nuclei were stained with DAPI. Bar = 10 μM. (**B**) Flow cytometry of Jurkat cells using an anti-TMPRSS13 (C-ter) antibody. Filled grey plot: non-labeled cells, gray line: isotype control, red line: TMPRSS13 staining. (**C**) Western blotting of TMPRSS13 expression in Derl, Myla, HuT78, SNK, Jurkat cells, purified CD3 lymphocytes, and purified B lymphocytes using anti-TMPRSS13 antibodies directed either to the C-terminal domain (C-ter), the entire extracellular domain, or the N-terminal domain (N-ter) of the protein. (**D**) Immunofluorescence surface labeling of PBMCs from healthy donors with anti-TMPRSS13 (green) and anti-CD3 (red) antibodies. Nuclei were stained with DAPI. Bar = 10μM. (**E & F**) Flow cytometry of TMPRSS13 of PBMCs labeled with Fixable Viability Dye and stained with anti-CD3-PE, anti-CD4-BV510, anti-CD8-BV711, anti CD14-FITC, anti-CD19-PC7, and anti-CD56-PC5.5. After fixation, cells were stained for TMPRSS13. Data were analyzed on the various populations after the exclusion of VD-positive cells. (**E**) Percentage of TMPRSS13-positive cells in each subpopulation. **(F)** MFI. n =6 donors. (**G**) Expression of TMPRSS13 by hMDM. Left: IF with anti-TMPRSS13 antibody of hMDM. Bar = 10 μM. Right: FCM of TMPRSS13 in hMDM. N=4.

### TMPRSS13 facilitates the processing and fusogenic properties of SARS-CoV2 spike

TMPRSS13 belongs to the same family of serine proteases as TMPRSS2, which cleaves SARS-CoV2 spike protein and facilitates envelope fusion with host cell membranes (*15*). TMPRSS13 is already known to facilitate the entry of influenza virus and coronavirus into host cells (*21*) and has been recently shown to cleave SARS-Cov2 spike in cells ectopically expressing TMPRSS13 (*17*). We first verified the capacity of TMPRSS13 to cleave the S1/S2 fragment of SARS-CoV viruses using a fluorogenic peptide cleavage assay with the HEK and HEK-T cell lines (**fig. 6A**). Indeed, the expression of TMPRSS13 allowed the cleavage of the SARS-CoV peptide (3.9-fold higher signal than in non-expressing HEK cells) and more efficiently that of the SARS-CoV2 peptide (7-fold increase). Digestion was blocked when the cells were pre-treated with Camostat, a known inhibitor of this family of serine proteases (*15*). To verify that TMPRSS13 expression facilitates the entry of SARS-Cov-2 LVS into cells, we transfected HEK and HEK-T cells with a plasmid encoding the primary virus receptor ACE2. The presence of TMPRSS13 facilitated the entry of GFP^+^ LVS into HEK-T cells relative to HEK cells, resulting in 40 ± 8.9% and 3.7 ± 2.1 % GFP^+^ cells at 48 hours post-infection, respectively (**fig. 6 B&C**). This corresponded to a more important processing of the spike protein into its fusogenic S2 fragment (**fig. 6D**). Human macrophages, which naturally express TMPRSS13, could be also infected by the spike pseudotyped virus (**fig. 6E**).

**Figure 6.**
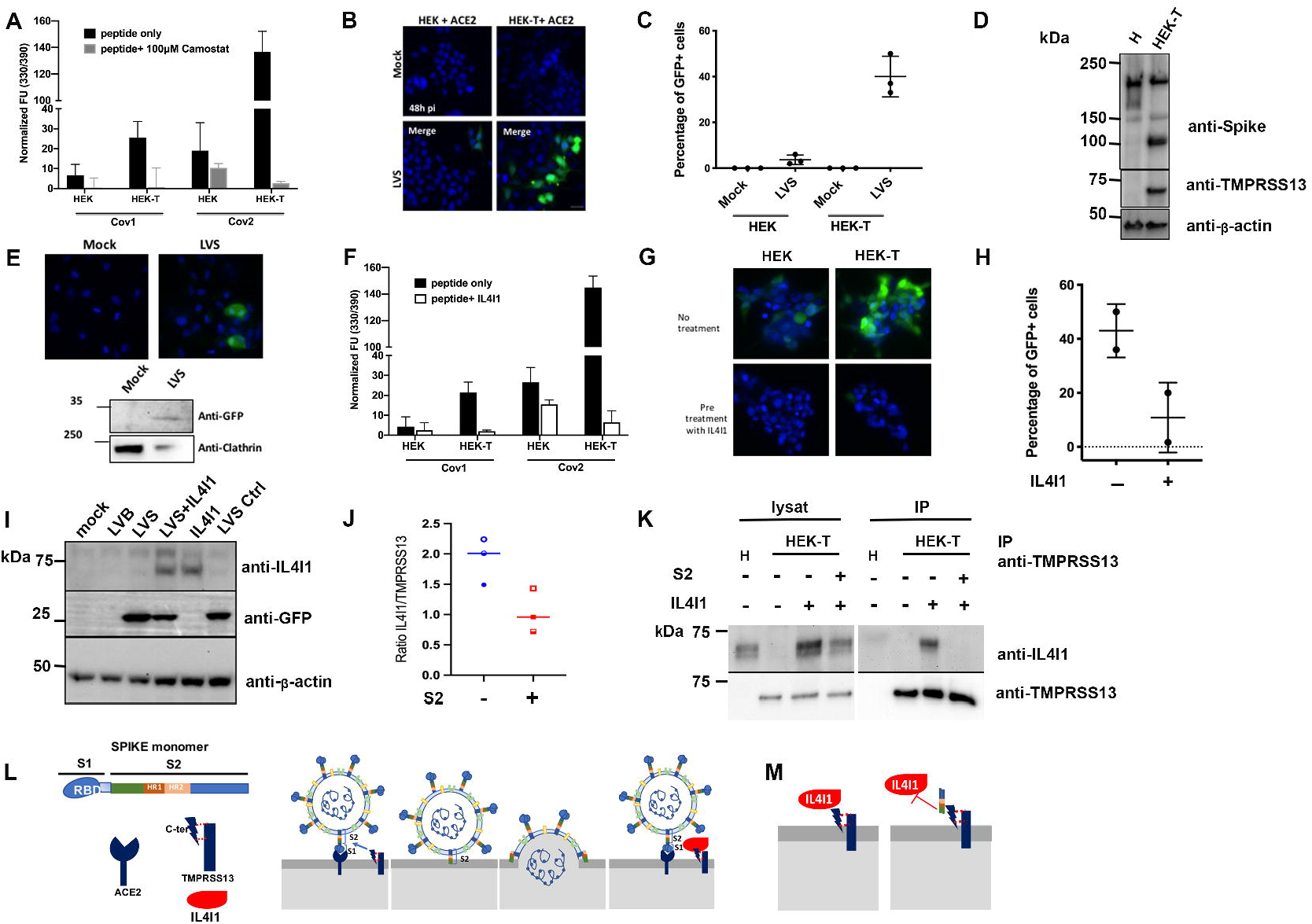
HEK cells expressing TMPRSS13 are infected by spike-pseudotyped virus and IL4I1 and spike compete for binding to TMPRSS13. **(A)** HEK or HEK-T cells were incubated at 37°C with either the fluorogenic peptide Cov1 (spike protein of the SARS-Cov) or Cov2 (spike protein of the SARS-Cov2). For Camostat treatment, cells were preincubated for 2 h with 1 mM Camostat, followed by the addition of 100 μm during the test. Fluorescence due to peptide cleavage was periodically measured. End-point measurements are expressed as arbitrary fluorescence units (UF) after subtraction of the background (fluorescence measurements without peptide). Data are presented as the average ± SD of 2 to 3 experiments, each performed in triplicate. (**B&C**) HEK and HEK-T cells were transfected with a plasmid expressing ACE2. Forty-eight hours later, cells were infected for 48 h with a spike pseudotyped virus (LVS) expressing GFP. Cells were observed under a fluorescent microscope (**B**) and GFP positive cells quantified (**C**). N =3. (**D**) HEK (H) or HEK-T (HT) cells transiently transfected for 48 h with a vector coding for spike protein were analyzed by WB using an anti-spike antibody followed by an anti-DYK antibody. Blots were rehybridized with an anti-actin antibody as a loading control. * S2 fragment. (**E**) hMDM were differentiated from monocytes for six days, infected with the LVS or mock infected for 48 h, and observed under a fluorescence microscope (top panels) or analyzed by WB (bottom panels). GFP (top blot) and clathrin used as loading controls (lower blot). Representative images are shown. (**F**) HEK or HEK-T cells as in (A), preincubated for 2 h with IL4I1. The test was performed as in (A). N=2-3. (**G**) HEK or HEK-T cells were pre-incubated or not with IL4I1 and infected with LVS and followed by IF. (**H)** Quantification of experiments shown in G. (**I**) hMDM pre-incubated or not with IL4I1 and infected with LVS for 48 h were analyzed by WB for IL4I1 (top panel), GFP (to detect LVS) (center panel), and □-actin (lower panel). As control, a bald pseudotyped virus (LVB) was used. A representative blot is shown. (**J & K)** HEK and HEK-T cells pre-incubated with 20 μg (J) or 80 μg (K) of recombinant S2 fragment for 2 h before IL4I1 addition were co-immunoprecipitated as in Fig. 4. (**J**) Quantified signals using ImageJ are expressed as the ratio of arbitrary units of IL4I1 and TMPRSS13. N=3. (**K**) Western-blot image of IL4I1/TMPRSS13 Co-IP performed in the presence of equivalent quantities (80 μg) of spike S2. (**L**) Schematic representation of the inhibition of the interaction of spike with TMPRSS13. Left: The spike protein interacts with ACE2 at the surface of target cells and TMPRSS13 cleaves the protein, liberating the S2 fragment, which changes conformation via the HR1 and HR2 domains, inserts into the host cell membrane, and favors envelope and cell membrane fusion. Right: if IL4I1 is present, it binds to TMPRSS13 and blocks it from interacting with the Spike protein and cleaving it. (**M**) Schematic representation of the inhibition of the interaction of IL4I1 with TMPRSS13. Left: IL4I1 binds to TMPRSS13 on the plasma membrane of T lymphocytes and inhibits their activation. Right: The spike S2 fragment binds to TMPRSS13 and prevents IL4I1 from binding to the lymphocyte, possibly blocking its effect.

### IL4I1 and SARS-CoV2 spike compete for the interaction with TMPRSS13

As both IL4I1 and spike interact with TMPRSS13, we looked for homology between these proteins. The two proteins show a small N-terminal region of homology, corresponding to their signal peptide. We found no homology before the S1/S2 cleavage site of spike. On the contrary, we found the S2 fragment to share moderate homology with IL4I1 (**fig. S4**), which reached 25% identity and 59% similarity in some regions. Most of the similarity resided in the S1/S2 cleavage region, Heptad Region (HR)1, HR2, and fusion domains of spike.

Considering this homology, IL4I1 may interfere with the ability of TMPRSS13 to cleave spike. We thus preincubated HEK-T cells with recombinant IL4I1 prior to testing cleavage of the S1/S2 peptide (**fig. 6F**). Indeed, IL4I1 almost completely prevented SARS-Cov2 peptide cleavage. Pre-incubation with recombinant IL4I1 also almost completely abolished the capacity of SARS-CoV2 pseudovirus to infect HEK-T cells (**fig. 6 G&H**). Moreover, the addition of IL4I1 to hMDM before infection with the SARS-CoV2 LVS significantly decreased entry of the virus (**fig. 6I**). Co-IPs of TMPRSS13 and IL4I1 in the presence of the S2 fragment of the spike protein (**fig. 6 J&K**) were next performed. When a large excess of IL4I1 (20 μg of S2 per 80 μg of IL4I1) was used, the presence of the S2 fragment decreased IL4I1 binding to TMPRSS13 by approximately 50% (**fig. 6J**). When the amount of S2 was equivalent to that of IL4I1, the binding of IL4I1 to HEK-T cells was completely abolished (**fig. 6K**). Thus, IL4I1 can interfere with the interaction between spike and TMPRSS13 and reciprocally, the spike protein interferes with IL4I1 binding to its receptor.

## Discussion

This study provides two major findings. First, we show the existence of a new receptor for the immunosuppressive enzyme IL4I1 and identify this protein as the serine protease TMPRSS13. This interaction may be important for IL4I1-mediated immunosuppression. Second, we show that TMPRSS13 facilitates spike cleavage and spike pseudotyped virus can enter immune cells. By interacting with TMPRSS13, IL4I1 and the spike protein can interfere with each other.

Although IL4I1 has several effects on the immune system, its mechanism of action remains partially elusive. In particular, the rapid effects on the TCR signaling pathway that have been observed (*5*) cannot be solely explained by the enzymatic activity of IL4I1, including the recently discovered AHR activation (*11*). The secretion of IL4I1 (*3*), and the observation that it can be found on the surface of T-cells after interaction with the cells that produce it (*5*), both suggested that IL4I1 can act as a cytokine. Here, using a combination of biochemical and immunological methods, we show that IL4I1 indeed binds to the transmembrane protein TMPRSS13, a serine protease recently shown expressed and important in human cancers and in viral infections (*21–23*). Its enzymatic activity combined with its acting through a receptor make IL4I1 a new “moonlighting” multitasking protein (*24*).

We show that TMPRSS13 is expressed by various immune cell types, with the percentage of positive cells greatly varying between donors, suggesting that the effect of IL4I1 could vary between individuals. Indeed, previously observed impact of IL4I1 on TCR signaling were donor dependent (*5*). The expression of TMPRSS13 by T and B lymphocytes may explain, at least partially, the action of IL4I1 on these cells. TMPRSS13 appears to be widely expressed by other circulating populations, suggesting that IL4I1 may have a larger spectrum of action in the immune system than currently known.

We observed that the C-terminal cleavage of TMPRSS13 is more prevalent and already present on primary T cells and cell lines where this protease may be already active. Such activation was not visible in primary B cells or in HEK-T cells overexpressing TMPRSS13. Serine transmembrane proteases show autocatalytic activity that is normally limited by Kunitz-type serine protease inhibitors, hepatocyte growth factor activator inhibitor (HAI)-1 or HAI-2 (*19*). T lymphocytes do not express these inhibitors and are often negative for other inhibitors, such as serpins (The Human Protein Atlas) (*25*). Their absence may explain the pre-activation of TMPRSS13 in T cells. The consequences of such activation are yet to be elucidated.

TMPRSS13 belongs to the same family of proteases as TMPRSS2, the protease involved in SARS-Cov2 spike protein cleavage (*15*). The role of TMPRSS13 in the processing of envelope proteins was already described for influenza virus and has been recently shown for coronavirus (*17*). We confirm that TMPRSS13 can facilitate the entry of SARS-CoV2 into cells that express it.

The interaction of IL4I1 and spike with TMPRSS13 led us to look for similarities between their sequences. Much to our surprise, the S2 fragment has several regions of similarity and/or homology with the human IL4I1 protein. These regions may be involved in the interaction of these proteins with TMPRSS13 and may indicate some convergent evolution (*26*).

The presence of IL4I1 blocked the digestion of spike and limited entry of the virus into the cells (see schema in fig. 6 L). This should be important for the effects of SARS-Cov2 on immune cells. Here, we show that macrophages can be infected by a pseudovirus bearing the spike protein. This is consistent with data from patients that indicate that both alveolar and monocyte-derived macrophages can be infected and modified by the virus (*27, 28*). As these cells are known to be resistant to viral infections but can harbor viruses, allow their replication, and behave as a Trojan horse (*29*), our observation could provide the basis for attractive anti-viral therapies to limit the consequences of immune cell infection.

Reciprocally, we have shown that the S2 fragment can block the interaction of IL4I1 with TMPRSS13 as depicted in fig.6 M. Such interference may be exploited to limit the immunosuppressive effect of IL4I1 in situations in which it is detrimental, such as cancer.

In conclusion, we have identified TMPRSS13 as the surface receptor for the immunosuppressive enzyme IL4I1 and have shown that IL4I1 and the envelope protein spike of SARS-CoV2 can interfere with each other for the interaction with TMPRSS13. Such interference open perspectives for the development of new treatments for COVID-19 or cancer patients.

## Acknowledgments

We thank the IMRB cytometry and imaging facilities as well the IMAG’IC platform (Institut Cochin) which is a member of the National Infrastructure France BioImaging (ANR-10-INBS04). The manuscript has been professionally corrected by a native English speaker from the scientific editing and translation company, Alex Edelman & Associates.

## Funding

This work was supported by:

Bristol-Myers Squibb Foundation for Research in Immuno-Oncology (FC)

Pre-maturation grant of Erganeo (FC).

INSERM, the CNRS, and Université de Paris (FN laboratory); INSERM and Université Paris Est-Créteil (UPEC) (JC laboratory)

## Author contributions

Designed experiments: FC, FN and VMF.

Performed experiments: JG, CN, AD, MPL, FN and FC

Analyzed the results: FC, FN, VMF, JG, JC and AD

Funding acquisition: FC, FN, VMF, JC

Wrote the manuscript - original draft: FC, VMF and FN

Wrote the manuscript – review & editing: FC, VMF, FN and JC

## Data and materials availability

anti-IL4I1 antibodies are available upon MTA. All data is available in the manuscript or the supplementary data.

