## Supplementary material for "IL4I1 binds to TMPRSS13 and competes with SARS-Cov2 Spike": Methods and supplementary figures: Supplementary Materials V2.pdf

Correspondence to:

### METHODS

#### Reagents

TriCEPS-biotin, TriCEPS-TAMRA Transferrin, and Na periodate were purchased from Dualsystems Biotech AG (Schlieren – Switzerland). The Pan T-cell isolation kit (130-096-535) and the B cell isolation kit II (130-091-151) were purchased from Miltenyi Biotech (Paris, France). Recombinant human IL4I1 (5684-AO) and spike S2 fragment (10594-CV) were purchased from R&D, Biotechne (Noyal-Chatillon-sur-Seiche, France). TMPRSS13 DNA (NM_001077263.2), angiotensin converting enzyme 2 (ACE2) DNA (NM_021804) and Anti-DYK antibodies (A00187) were purchased from Genscript (Leiden, Netherlands). Anti-TMPRSS13 polyclonal antibodies (AP14675b) and Protein A/G agarose beads (Sc2003) of Santa Cruz Biotechnology were purchased from Cliniscience (Nanterre, France). The anti-TMPRSS13 N-terminal antibody (Ab59862) was purchased from Abcam (Paris, France). All culture reagents were purchased from Life Technologies. Anti-TMPRSS13 antibody (Pa5-30935), Texas-red phalloidin (T7471), goat anti-rabbit-Alexa Flour 488 (A21206), goat anti-rabbit AlexaFlour 647 (A21244) antibodies, poly D-lysine (A389040), ProLong™ Gold Antifade Mounting Media (P10144), Lipofectamine 2000 (11668027), eBioscience Fixable Viability Dye-efluor 450 (65-0863), anti-human CD4 APC-eFlour 780 (47-0049-42, clone RPA-T4), anti-human CD28 (16-0289-81, clone 28.2), and their respective isotype controls were purchased from ThermoFisher (Villebon-sur-Yvette, France). Anti-human CD14-FITC (555397, clone M5E2), anti-human CD3-PE (555333, clone UCHT1), and their respective isotype controls were purchased from BD Biosciences (Pont de Claix, France). Anti-human CD56-PC5.5 (A79388, clone N901), anti-human CD19-PC7 (IM3628, clone J3-119), and their respective isotype controls were purchased from Beckman Coulter. Anti-human CD4-BV510 (BLE300546, clone RPA-T4), anti-human CD8 BV711 (BLE301044, clone RPA-T8), their respective isotype controls, True Staining Monocyte blocker (42610), and PE-streptavidin (405203) were purchased from Biolegend (Amsterdam, Netherland). Goat-anti-rabbit-HRP (7074) and horse-anti-mouse-HRP (7076) antibodies from Cell Signaling were purchased from Ozyme (Saint-Cyr-L'École, France). Pre-cast SDS-PAGE gels, Precision Plus Protein™ Dual Color Standards (1610374) and TidyBlot Western Blot Detection Reagent-HRP (STAR209) were purchased from BioRad (Marne-la-Coquette, France). Gel running buffer (TG-SDS EU0510) and transfer buffer (TG EU0550) were purchased from Euromedex (Souffelweyersheim, France). Roche cOmplete™ Mini Protease Inhibitor Cocktail (4693124001), anti-myc monoclonal antibodies (M4439, clone 9E10), Millipore Immobilon-P

PVDF membranes (IPVH00010), Millipore Luminata Crescendo, Amicon ultra 4 30K (UFC803008), and the Duolink PLA kit (DU092007) were purchased from Merck Millipore (Guyancourt, France). Monoclonal anti-rabbit antibodies against IL4I1 were produced against a peptide by our laboratory (clone 43-7, patent EP18306563.0). Concentrated 32% Paraformaldehyde (PFA) was purchased from Electron Microscopy Sciences (Biovalley, France) and diluted in PBS at a final concentration of 4%.

**Cells**

Jurkat cells were cultivated in RPM1 1640 containing 10% fetal calf serum (FCS), 100 units/ml penicillin, and 100 μg/ml streptomycin. Human embryonic Kidney (HEK) cells were cultivated in DMEM containing 10% FCS, 100 units/ml penicillin, and 100 μg/ml streptomycin. HEK cells expressing inducible recombinant human IL4I1 (rhIL4I1) have been described elsewhere (*3*) and were used in certain experiments (PLA and co-IP). For these experiments, IL4I1 was induced using 2 mg/ml doxycycline for a week from sub-confluent cells. Over the last 16 h, the recombinant protein was recovered in PBS containing $Ca^{++}$ and $Mg^{++}$ and subsequently concentrated using an Amicon filter with a cut-off of 30 kDa. After concentration, the IL4I1 activity and protein content were measured. A volume of the conditioned PBS corresponding to 10,000 U of specific activity (pmol $H_2O_2$/h/mL IL4I1) was added to a well containing the cells in a six-well plate.

Peripheral blood mononuclear cells (PBMCs) were obtained by cytapheresis from healthy donors from the French Blood bank (Etablissement Français du sang, EFS) and isolated using a Ficoll density gradient. Monocyte-derived macrophages (hMDMs) were differentiated from PBMCs according to the method described in Lê-Bury et al. (*30*) for six days. $CD3^+$ cells and naïve B cells were isolated from donor PBMCs using the magnetic bead Pan T isolation kit and B Cell Isolation Kit II from Miltenyi respectively, according to the manufacturer's instructions.

**Pseudovirus**

EGFP-expressing pseudovirus (LVS) based on a pLV[Exp]-CMV>EGFP lentivirus and typed with spike protein or bald controls (LVB) were purchased from Vectorbuilder Gmb. For the infections, cells plated in six-well plates (transfected HEK or hMDM) were washed with PBS followed by the addition of 500 µL medium without antibiotics containing the diluted virus and incubation for 1 h at room temperature out under gentle agitation. Plates were washed with medium and fresh complete medium was added for the rest of the experiment.

### DNA

The sequence of TMPRSS13 (NM_001077263.2) was cloned into a pcDNA3.1(+)-C-DYK vector and that of ACE2 (NM_021804) into a pcDNA3.1(+)-C-HA vector. All DNA constructs were produced by Genscript (Leiden, Netherland).

### Transfections

Transfection of HEK cells with TMPRSS13 DNA was performed using Lipofectamine 2000 according to the manufacturer's instructions. For stable transfectants, cells were diluted the day after transfection and the following day selection started using 1.2 mg/mL G418. The selection media was changed every two days until the appearance of antibiotic-resistant cell colonies. Each colony was then isolated and verified for protein expression by WB. One clone was chosen for the rest of the experiments (HEK-T6).

### TriCEPS experiments

Ligand receptor-based capture (LRC) and mass spectrometry was performed using LRC-TriCEPS-biotin (Dualsystems Biotech), as previously described (*18*). Briefly, capture was performed on human $CD3^+$ T cells or Jurkat cells using hrIL4I1 bound to the LRC-TriCEPS bait. Anti-human CD28 antibody or transferrin were used as positive control baits and glycine as a negative control. For LRC-TriCEPS binding, 240 µg of control bait or rhIL4I1 protein were buffer exchanged to 150 µl 25 mM HEPES, pH 6.5. LRC-TriCEPS V.3 was added to each reaction and the reaction mixed and incubated at 22°C with gentle shaking for 90 min. After the reaction, the bound baits were quenched for 30 min on ice with an equal volume of a 125 mM Tris-HCl solution, pH 6.5.

For FCM binding experiments, 0.5 x $10^6$ cells (Jurkat or purified $CD3^+$ lymphocytes) were labeled with Fixable Viability Dye efluor 450 for 30 min at room temperature. Cells were then incubated with 1.2 µg anti-CD28 antibodies ($CD3^+$ cells), transferrin (Jurkat cells), or hIL4I1 previously bound to LRC-TriCEPS (TriCEPS-conjugated) for 1 h on ice. After extensive washing, LRC-TriCEPS-baits bound to cells were stained with streptavidin-PE and CD4 positive lymphocytes identified using an anti-CD4-APC-eFlour 780 antibody and analyzed on a LSRII flow cytometer (BD Bioscience, France). Analysis was performed using Flowjo software.

For mass spectrometry identification of interacting proteins, 1.2 x $10^8$ Jurkat cells were oxidized by treatment with 1.5 mM sodium metaperiodate at 4°C for 15 min. After oxidation, the cells were washed, divided into two parts, and incubated with TriCEPS-transferrin or TriCEPS-rhIL4I1 (see above) for 90 min in the dark under rotation. For each condition, cells were divided into three tubes and frozen at -80°C. Mass spectrometry analysis was performed using the Dualsystem approach according to Frei et al. . Samples were analyzed on a Thermo LTQ Orbitrap XL spectrometer fitted with an electrospray ion source. Tryptic peptides were measured in data dependent acquisition mode (TOPN) in a 120 min gradient using a 10-cm C18 packed column. Progenesis software was used for raw file alignment and feature detection, the Comet Search Engine for spectra identification, and the Trans-Proteomic Pipeline for statistical validation of putative identifications and protein inference.

Upon protein inference, relative quantification of the control and ligand samples was performed based on the ion extracted intensity and differential protein abundance was tested using a statistical ANOVA model, followed by a multiple testing correction. This model assumes that the measurement error follows a Gaussian distribution and views individual features as replicates of a protein's abundance and explicitly accounts for such redundancy. It tests each protein for differential abundance in all pairwise comparisons of ligand and control samples and reports the p-values. The Uniprot human proteome data base was used for analysis.

For TAMRA-TriCEPS experiments, Jurkat cells were incubated with TAMRA-TriCEPS Transferrin or TAMRA-TriCEPS-rhIL4I1 for 1 h on ice in the dark. Following incubation, cells were washed and resuspended in PBS containing 0.1% FCS and 6.5 x $10^4$ were spun onto 12-mm glass coverslips previously coated with 0.1% poly-D-lysine. Cells were then fixed with 4% para-formaldehyde (PFA) for 20 min and the coverslips mounted with Prolong containing DAPI. Cells were observed using a fluorescence Axioimager M2 EC (Zeiss, France) with Plan Neofluar 40X/0.75 and x63/1.25 objectives.

**Immunofluorescence**

HEK cells were grown on glass coverslips pre-coated with 0.1% poly-D-lysine. hMDMs were cultivated and differentiated on glass coverslips. Cells were fixed with 4% PFA for 20 min at RT. For hMDMs, 1 µL of True Stain monocyte blocker was added for 10 min before labeling to each coverslip to block FC receptors. For non-permeabilized cells, coverslips were directly blocked with 10% bovine serum albumin (BSA) followed by incubation with primary and secondary antibodies. For permeabilized cells, the cells were incubated for 4 min at room temperature in a 0.1% Triton X100 solution before being blocked with BSA. After staining, coverslips were mounted using Prolong containing DAPI and observed using a fluorescence Axioimager M2 EC using Plan Neofluar 40X/0.75 and x63/1.25 objectives.

**Proximity ligation assay (PLA)**

PLA was performed according to the manufacturer's instructions. Briefly, HEK or HEK-T expressing TMPRSS13 were plated onto 0.1% poly-D-lysine-treated coverslips in 24-well plates the day before the assay. Cells were incubated for 1 h on ice with conditioned PBS containing rhIL4I. After extensive washing with PBS, cells were fixed with 4% PFA for 20 min, blocked, and incubated with mouse monoclonal anti-DYK and rabbit monoclonal anti-IL4I1 antibodies for 30 min at 37°C. Cells were then incubated with the secondary antibodies bound to the oligonucleotide probes for 1 h at 37°C. Following several washes, a solution containing DNA ligase was added to the samples, followed by a 30 min incubation at 37°C. Finally, washed samples were incubated in an amplification solution containing polymerase and a Cy3 fluorescent nucleotide. Coverslips were then washed and mounted with Prolong containing DAPI and observed under a fluorescence microscope with a 20X Acroplan x20/0.45 objective. For PLA experiments with Jurkat cells, mouse monoclonal anti-myc and rabbit polyclonal anti-TMPRSS13 antibodies were used.

**Co-immunoprecipitation**

HEK or HEK-T cells from a six-well plate were incubated for 1 h on ice with concentrated hIL4I (10000U/well). After 10 washes with 1 ml PBS, cells were lysed in 500 µL lysis buffer (50 mM Tris HCl, pH 7.5, 150 mM NaCl, 2 mM EDTA, 0.5% Triton X100) containing cOmplete™ Protease Inhibitor Cocktail. Lysates were spun at 5,000 x g for 5 min and whole cell supernatants incubated overnight with rotation at 4°C with 1.5 µg anti-DYK antibodies or 3 µg anti-IL4I1 antibodies (clone 43-7, patent EP18306563). After the addition of 50 µL of a 50% suspension of protein A/G agarose beads and incubation for an additional 2 hat 4°C, the agarose beads were washed three times with 1 mL lysis buffer and resuspended in 50 µL Laemmli sample buffer (0.1 M Tris-HCl, pH 6.8, 10% glycerol, 1% SDS, 0.05 M DTT, 0.2% bromophenol blue). After an incubation of 5 min at 95°C and samples were loaded onto a 10% SDS-PAGE. Ten microliters of the whole-cell supernatants were run in parallel as input controls. For the experiments with the S2 fragment of the spike protein, 20 or 80 µg of the recombinant protein were added on the cultures 2 h before the addition of IL4I1 as described above.

**Flow cytometry**

Viable PBMCs were identified using Fixable Viability Dye efluor 450. Then, surface labeling was performed (anti-human CD3-PE, CD4-BV510, CD8-BV711, CD14-FITC, CD19-PC7, and CD56-PC5.5) by adding the antibody cocktail containing True Stain monocyte blocker on ice and performing all the steps at 4°C. After washing, PBMCs were fixed with 4% PFA. For Jurkat cells and HEK-T cells, the cells were directly fixed. The fixed cells were washed in PBS and resuspended in PBS containing 1% FCS and 10 ng/ml of the anti-TMPRSS13 antibody and incubated for 30 min at 4°C, followed by incubation with goat ant-rabbit Alexa Fluor 647 antibody. Data was collected using a Fortessa X20 flow cytometer (BD Bioscience, France) and analyzed using FlowJo software.

**Western blots**

Proteins separated by SDS-PAGE (pre-cast gels NuPAGE Themo Scientific or BioRad) were transferred to PVDF membranes (Merck Millipore, France) membranes at 100 mV for 2 h or overnight at 30 mV in 25 mM Tris, 192 mM glycine, and 20% (v/v) ethanol. Membranes were blocked with a 10% BSA solution in TBS-T (50 mM Tris-Cl, pH 7.6, 150 mM NaCl, 0.05% Tween-20) before incubation with specific antibodies diluted in 1% BSA-TBST. Primary antibodies were revealed using anti-rabbit-HRP, anti-mouse-HRP antibodies or TidyBlot Western Blot Detection Reagent-HRP using Luminata Crescendo (Merck-Millipore, France) or Pierce Dura. Blots were re-probed after HRP inactivation by incubation in 30% $H_2O_2$ (8.8 M) for 30 min at 37°C according to Sennepin et al. (*31*). Images were captured using a CCD camera (Autochemi system, UVP, UK) or Fusion FX devices (Vilber, France) and analyzed using ImageJ.

**IL4I1 activity**

IL4I1 LAAO activity was measured against Phe as previously described (*7*).

**Spike in-vitro digestion system**

The digestion test was performed according to Jaimes et al. (*32*). Peptides corresponding to the SARS-CoV and SARS-CoV-2 spike (S) S1/S2 sites composed of the sequences HTVSLLRSTSQ and TNSPRRARSVA, respectively, and harboring the (7-methoxycoumarin-4-yl)acetyl/2,4-dinitrophenyl (MCA/DNP) FRET pair were synthesized by Biomatik (Wilmington, DE, USA). Briefly, HEK and HEK-T cells were seeded at $10^4$ cells/well in a flat bottom 96-well plate the day before the experiment. Cells were washed with 200 µL PBS and then incubated with 200 µM of each peptide. For certain experiments, cells were preincubated for 2 h with 1 mM Camostat or 1500U/well concentrated recombinant IL4I1. Camostat was maintained at 100 µM during the assay. Plates were placed in a Varioskan™ LUX plate reader and fluorescence emission at 37°C was kinetically recorded using λex 330 nm and λem390 nm wavelengths for up to 45 min. Data are shown as the mean values for the endpoints of triplicates from which the background values (fluorescence of PBS only) were subtracted.

**Protein alignment**

The Spike (sp|P0DTC2|SPIKE_SARS2) and IL4I1 protein sequences (NP_690863.1) were aligned using Clustal Omega and analyzed using Jalview (*33*).

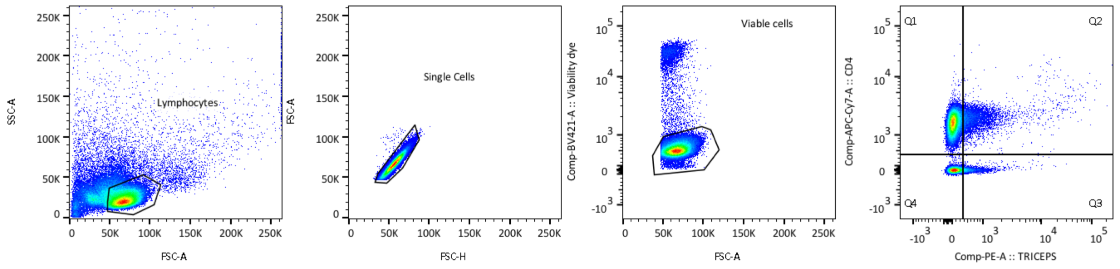


**Fig. S1**. Gating strategy for TriCEPS labeling. Lymphocytes were identified on the SSC-A FSC-A plot. After the exclusion of doublets, cells were gated on Viability Dye (VD)-negative cells. For Jurkat cells, the LCR-TriCEPS labeling was measured in this window. For lymphocytes, the LCR-TriCEPS labeling a was further analyzed on the CD4+ population.

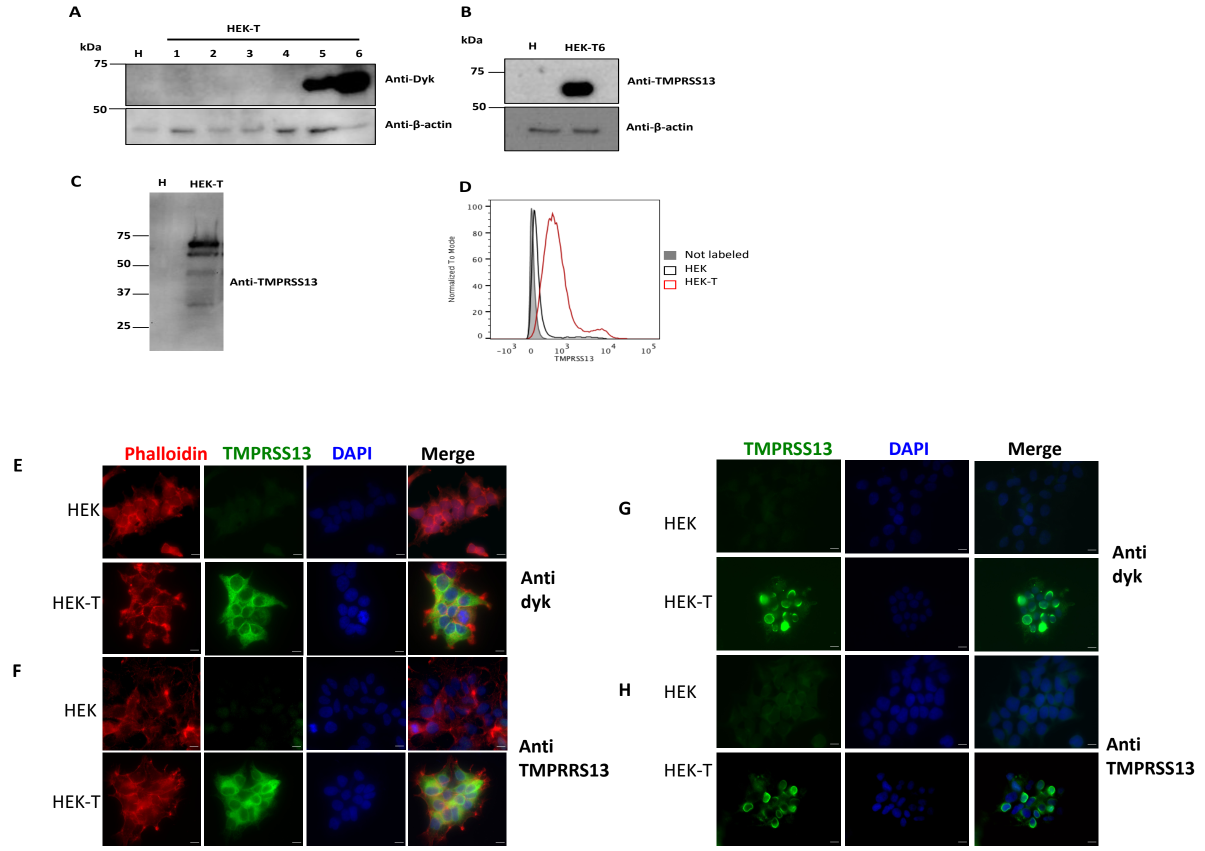


**Fig. S2. Validation of the HEK clone expressing TMPRSS13 and the commercial anti-TMPRSS13 antibody.** HEK cells stably transfected with human DYK-tagged TMPRSS13 cDNA were analyzed by WB using an anti-DYK antibody (**A**). HEK clone HT6, expressing TMPRSS13, was selected for further studies. Clone HEK-T6 (**B**) and transiently transfected cells (**C**) were tested using a commercially available anti-TMPRSS13 antibody directed against the C-terminal part of TMPRSS13 by WB. (**D**) HEK cells and HEK cells overexpressing TMPRSS13 (HEK-T) were tested by FCM using the anti-TMPRSS13 antibody. Filled grey plot: non-labelled cells, black line: HEK cells, red line: HEK-T cells. **(E)** Triton X100 permeabilized HEK (top) and HEK-T (bottom) cells were labeled with an anti-DYK antibody followed by an anti-mouse Alexa488 antibody. **(F)** Triton X100 permeabilized HEK (top) and HEK-T (bottom) cells were labeled with an anti-TMPRSS13 rabbit polyclonal antibody, followed by an anti-rabbit Alexa 488 antibody. Texas red phalloidin: actin cytoskeleton and DAPI: nuclei. Bar = 10 µM. (**G**) Non-permeabilized HEK (top) and HEK-T cells (bottom) were labeled with an anti-DYK antibody, followed by an anti-mouse Alexa 488 antibody. (**H**) Non-permeabilized HEK (top) and HEK-T cells (bottom) were labeled with an anti-TMPRSS13 antibody, followed by an anti-rabbit Alexa 488 antibody. DAPI : nuclei. Bar = 10 µM.

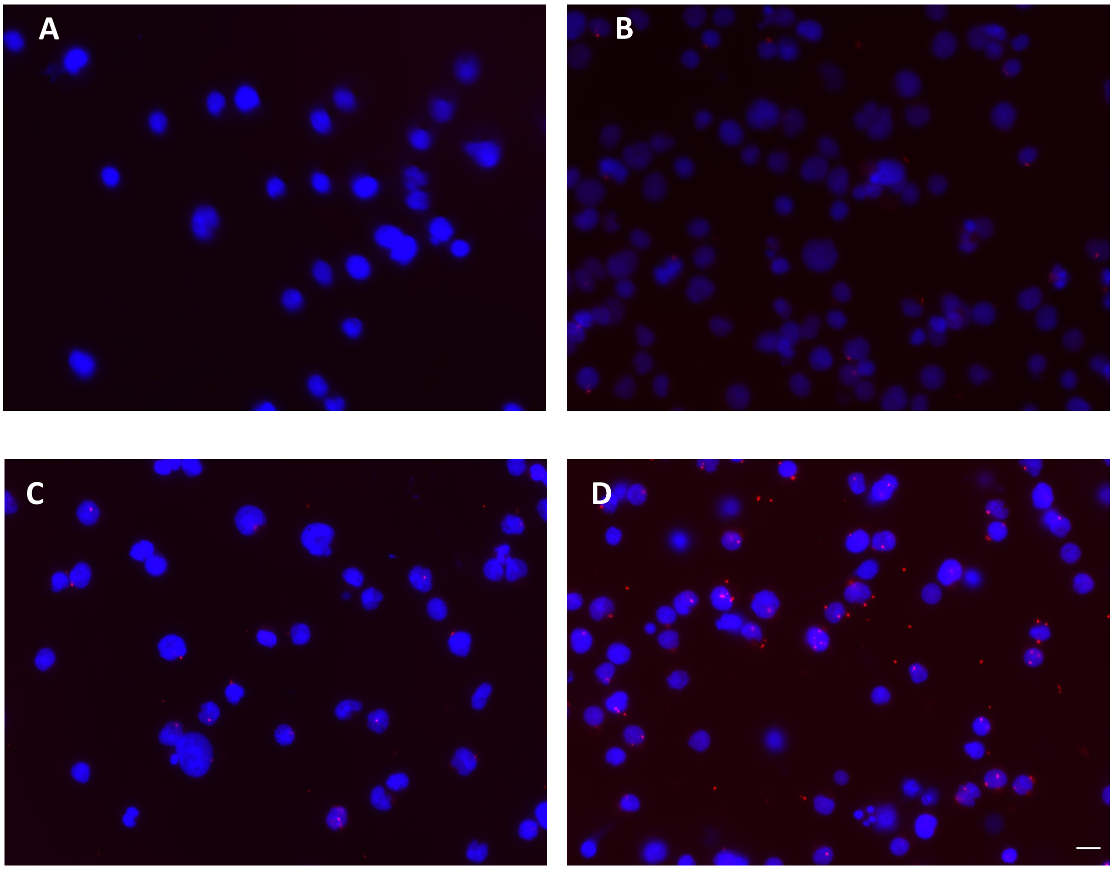


**Fig. S3. Proximity ligation assay of Jurkat cells incubated reveals spatial proximity of IL4I1 and TMPRSS13**. Jurkat cells were incubated with recombinant hIL4I1 at 4°C for 1 h. After extensive washing and 4% PFA fixation, cells were incubated with rabbit anti-TMPRSS13 and/or mouse anti-myc primary antibodies and secondary anti-mouse and anti-rabbit antibodies bound to specific primers. After ligation and the PCR reaction containing a red fluorescent nucleotide, the coverslips were mounted with DAPI to stain the nuclei. (**A**) No primary antibodies, (**B**) primary anti-DYK antibody only, (**C**) Jurkat cells without incubation with IL4I1 or primary or secondary antibodies, **(D**) Jurkat cells incubated with IL4I1 and the primary and secondary antibodies. Images were captured using an Axioimager 2 fluorescent microscope and analyzed using ImageJ. Representative images from three independent experiments (n = 3). Bar = 10 μM.

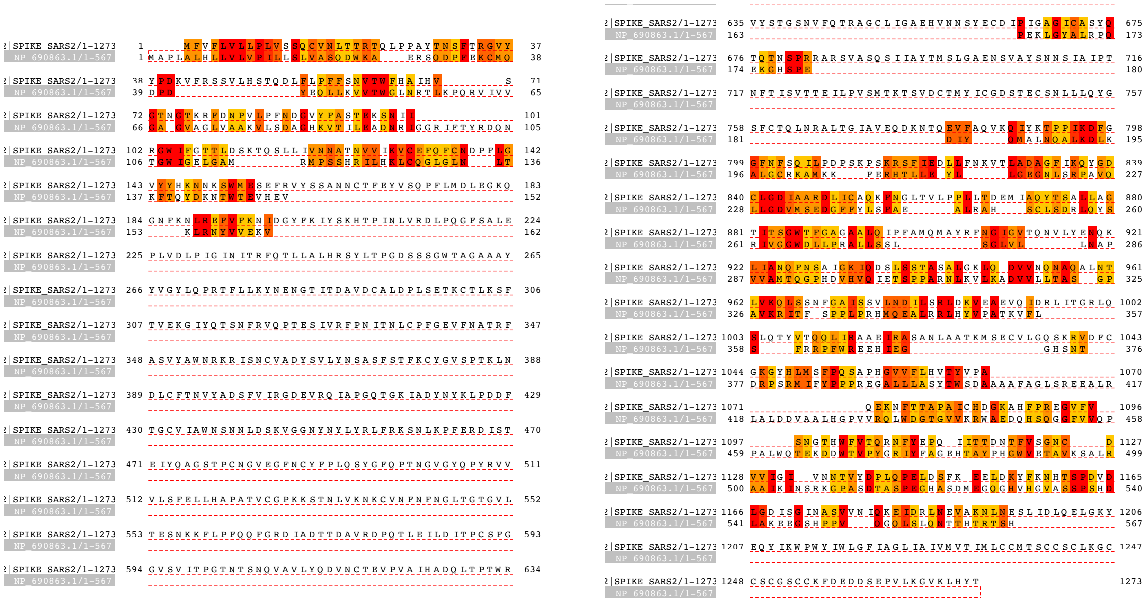

**Fig. S4. Protein alignment of Spike and IL4I1**. The Spike (sp|P0DTC2|SPIKE_SARS2) and IL4I1 protein sequence (NP_690863.1) were aligned using Clustal Omega and analyzed using Jalview. Homologous and similar amino acids are annotated in shades from red to yellow (Threshold 6.996).
